# Non-crossover gene conversions show strong GC bias and unexpected clustering in humans

**DOI:** 10.1101/009175

**Authors:** Amy L. Williams, Giulio Genovese, Thomas Dyer, Nicolas Altemose, Katherine Truax, Goo Jun, Nick Patterson, Simon R. Myers, Joanne E. Curran, Ravi Duggirala, John Blangero, David Reich, Molly Przeworski, T2D-GENES Consortium

## Abstract

Although the past decade has seen tremendous progress in our understanding of fine-scale recombination, little is known about non-crossover (NCO) gene conversion. We report the first genome-wide study of NCO events in humans. Using SNP array data from 98 meioses, we identified 103 sites affected by NCO, of which 50/52 were confirmed in sequence data. Overlap with double strand break (DSB) hotspots indicates that the events are likely of meiotic origin. We estimate that a site is involved in a NCO at a rate of 5.7×10^-6^/bp/generation, consistent with sperm-typing studies, and infer that tract lengths span at least an order of magnitude. Observed NCO events show strong allelic bias at heterozygous AT/GC SNPs, with 68% (58–78%) transmitting GC alleles (*P*=5×10^-4^). Strikingly, in 4 of 15 regions for which there are also resequencing data, multiple disjoint NCO tracts cluster in close proximity (~20–30 kb), a phenomenon not previously seen in mammals.

## Introduction

Meiotic recombination is a process that deliberately inflicts double strand breaks (DSBs) on the genome, leading to their repair as either crossover (CO) or non-crossover (NCO) resolutions. COs play an essential role in the segregation of chromosomes during meiosis whereas NCOs are thought to aid in homolog pairing or in shaping the distribution of COs over the genome [1,2]. While the past decade has seen tremendous progress in our characterization of DSBs and COs in mammals [1], little is known about NCO events.

These two resolutions appear to result from a choice early on in the repair of DSB breaks [3], with a number of properties differing between them [2,4]. In particular, both outcomes are accompanied by a short gene conversion tract that fills in the DSB on one homologous chromosome with the sequence from the other homolog. Whereas COs yield chromosomes with multi-megabase long segments from each homolog [1], NCO gene conversion tracts have been estimated to span ~50–1,000 bp [5]. Although short, these NCO gene conversion tracts affect sequence variation by breaking down linkage disequilibrium (LD) within a localized region, and, together with COs, are necessary to explain present-day haplotype diversity [6,7].

Despite the importance of NCOs, the frequency with which they occur in mammals remains uncharacterized. Estimates based on the number of DSBs that occur in meiosis suggest that NCOs are an order of magnitude more frequent than COs [8,9]. In turn, sperm-typing studies and analyses of LD indicate that NCOs occur ~1 to 15 fold more frequently than COs [5–7,10,11], with this value varying widely in analyses of individual hotspots [5,11]. Furthermore, while COs occur at a higher rate in females than in males and tend to occur in different genomic locations [12], there has been no comparison between the sexes for NCO events.

The locations of NCO events with respect to recombination hotspots is of interest more broadly. While NCO events are assumed to occur at the same hotspots for DSBs as COs [1], in humans, this has only been demonstrated for a limited number of locations in sperm [13]. Furthermore, by considering events in a single meiosis, sperm-typing studies have identified *complex crossovers* in which gene conversions tracts occur near but not contiguous with CO breakpoints [14]. A genome-wide analysis of NCO may therefore reveal further features of recombination.

The impact of NCO events on genome evolution is also in need of quantification. Cross-species analyses have shown that in highly recombining regions, GC content increases over evolutionary time, consistent with an important role for GC-biased gene conversion (gBGC) [15]. Polymorphism data also reveal an effect of recombination, with more AT to GC polymorphisms observed in regions of higher recombination [16,17]. Moreover, because gBGC acts analogously to positive selection, its effects on polymorphism and divergence can confound studies of human adaptation [18]. Although one recent sperm-typing study reported two recombination hotspots that exhibit GC-bias in NCO resolutions [11], most of the evidence of gBGC in mammals is indirect.

Motivated by these considerations, we carried out a study of NCO gene conversion events in pedigrees—to our knowledge, the first genome-wide assay of *de novo* NCO gene conversion in mammals. We sought answers to the following questions: (1) Do NCO events localize to the same hotspots as COs? (2) What is the rate at which a site is a part of a NCO tract? This is equivalent to the fraction of the genome affected by NCO in a given meiosis. (3) Are there differences in the NCO rate or localization patterns between males and females? (4) What is the strength of NCO-associated gBGC across the genome? (5) Do NCO gene conversion tracts vary substantially in length? (6) Do complex resolutions occur, with discontinuous tracts within a short distance?

We utilized two different sources of data for our analysis. The primary analysis focused on SNP array data from 34 three-generation pedigrees. These SNP array data provide information from 98 meioses, 49 paternal and 49 maternal, and are informative at 12.1 million sites (markers where we can potentially detect a NCO in a parent-child transmission). We followed up with a secondary analysis of a subset of the identified NCO events using whole genome sequence data.

## Results

We carried out a study of *de novo* meiotic NCO gene conversion resolutions in humans by analyzing Illumina SNP array data at two SNP densities (660k and 1M SNP density arrays; see Methods “Samples and sample selection”) from 34 three-generation Mexican American pedigrees [19–21]. The goal was to identify *de novo* NCO gene conversion events, manifested as one or more adjacent SNPs at which the alleles descend from the opposite haplotype relative to flanking markers (Figure 1a). Identifying these NCO events requires phasing of genotypes in the pedigree in order to infer haplotypes and the locations of switches between parental homologs in transmitted haplotypes.

**Figure 1.**
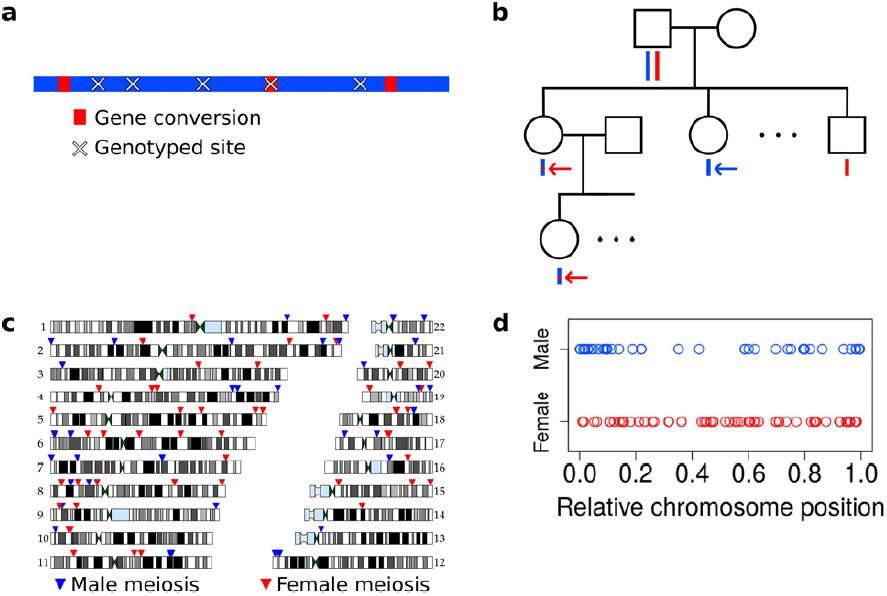
Non-crossover detection. **a**, Pictorial representation of a haplotype transmission including NCO events. A parent has two copies of each chromosome but transmits only one copy to his or her children. That copy is composed of DNA segments from the parent’s two homologs; i. e., it is formed by recombination between these two haplotypes. Here, the two haplotypes in the parent are colored in blue and red, and switches in color represent sites of recombination. The figure only depicts short NCO events and no COs. Overlaid on this haplotype are × symbols representing sites assayed by the SNP array. In this example, only one NCO has a SNP array site within it and only that NCO can be identified. **b**, To avoid calling false positive NCO events driven by genotyping error, we required putative NCO events first to be detected in a second generation child (top red arrow) and also transmitted to a third generation grandchild (bottom red arrow). We also required that the allele from the opposite haplotype (i.e., the one not affected by the NCO) in the parent (first generation) be transmitted to at least one child in the second generation (blue arrow). This study design ensures that false positive NCOs will only occur if there are two or more genotyping errors at a site. All 34 pedigrees included in this study have genotype data for both parents, at least three children, one or more grandchild, and both parents of included grandchildren. **c**, Genomic locations of the NCO sites that we detected are indicated by arrowheads, with red arrowheads representing NCO events from female meioses, and blue from male meioses. Many of the male NCO events localize to the telomeres. **d**, Relative chromosomal positions of events, stratified by the sex of the transmitting parent.

Two features make locating NCO events challenging. The first is the density of informative sites. NCO gene conversions have an estimated mean tract length of 300 bp or less [5,11], but on a SNP array with ~1 million variants, genotyped sites occur on average every 3,000 bp. Thus SNP array data will identify only a small subset of NCO events. Moreover, to be informative about NCO events (and recombination in general), a site must be heterozygous in the transmitting parent, so not all assayed positions are informative.

The second challenge arises from erroneous genotype calls. Errors in SNP array data can in principle confound an analysis of NCO because certain classes of errors can mimic these events (e.g., if a child is truly heterozygous but is called homozygous, or if a parent is homozygous but called heterozygous). Our study design minimizes false positive NCO calls by using three-generation pedigrees, as depicted in Figure 1b. The approach requires that a putative NCO event identified in a child in the second generation is also transmitted to a grandchild (red arrows in Figure 1b). Additionally, the approach validates the genotype of the transmitting parent as heterozygous by requiring that the allele from the alternate haplotype in that parent (i.e., the one that is not affected by NCO) be transmitted to at least one child (blue arrow in Figure 1b). These requirements exclude the possibility that a segregating deletion will be misinterpreted as a NCO event. Moreover, they guarantee that a false positive NCO event will only be called if there are at least two genotyping errors at a site. Specifically, for a false positive to occur, either the recipient of the NCO and his or her child must be incorrectly typed, or the parent transmitting the putative NCO and the child/children receiving the alternate allele must be in error. This approach decreases the number of events that can be detected since not all sites affected by NCO will be transmitted to a grandchild, but importantly it also greatly reduces the false positive rate. Further details on data quality control measures appear in Methods (“Quality control procedures” and “Pedigree-specific quality control”).

Our approach for identifying NCO events consisted of, first, phasing each three-generation pedigree using the program HAPI [22] (Methods “Phasing”). Next, we identified informative sites relative to each parent in the first generation: sites where the parent is heterozygous, the inferred phase is unambiguous, and where, if a NCO event occurred, both alleles are seen transmitted to children (see Methods “Determination of informative sites”). We then examined all apparent double CO events that occur within a span of 20 informative sites or less, i.e., we identified haplotype transmissions that contain switches from one parental haplotype to the other and then switch back to the original haplotype. Most of these recombination intervals span one to three SNPs and are less than 5 kb; these are putative NCO events. A few loci showed complex patterns with multiple, discontinuous recombination events across several SNPs, with tracts spanning 5 kb or more; these are not counted as NCOs but are described further below.

We ascertained the total number of informative sites in the same way as our NCO events. Thus, when calculating the per base pair (bp) rate of NCO, the numerator and denominator are identically ascertained (see below and Methods “Determination of informative sites” for details).

### Inferred non-crossovers and their likely source

Within the 34 three-generation pedigrees, we considered transmissions from a total of 98 first generation meioses (49 paternal, 49 maternal). This analysis revealed a total of 103 SNP sites (henceforth “NCO sites”) putatively affected by autosomal NCO events: 97 with standard ascertainment, and an additional six that are detectable but do not meet all the criteria for inclusion in the rate calculation (Figure 1c; Supplementary file 1; Methods “Determination of informative sites”). Most (76/103) NCO events derive from a single SNP, while others contain two or three NCO sites that delimit a tract. The NCO sites have roughly equal numbers of homozygous and heterozygous genotype calls in the recipient (56% heterozygous sites, *P*=0.59, two-sided binomial test), as expected, providing further confidence that the calls are not spurious. Furthermore, we confirmed genotype calls for a subset of the putative NCO events using whole genome sequence data generated by the T2D-GENES Consortium. Sequence data were available for 52 of these NCO sites, of which 50 are concordant with the SNP array calls (Methods “Validating non-crossover events”, Supplementary file 1). Of the two discordant sites, one shows evidence of being an artifact in the sequence data rather than the SNP array data, and for the other, the source of error is unclear (see Methods “Validating non-crossover events”). Overall, the error rates in these data are low, and so in what follows we assume that all 103 detected NCO events are real.

Meiotic NCOs are thought to localize to the same hotspots as COs [1], and studies at specific loci in sperm have supported this hypothesis [13]. To evaluate this question using genome-wide data, we utilized CO rates that Kong *et al.* estimated based on events identified in an Icelandic pedigree dataset [12]. This genetic map omits telomeres, and thus these rates are only available for a subset of our identified NCOs. The overlapping *de novo* NCOs show strong enrichment in sites with sex-averaged CO rate ≥10 cM/Mb (Figure S1). Indeed, 18 of the 72 events that we can examine (26%) localize to such regions (using only one SNP per NCO event), while 4.2% of informative sites have this high a rate. This co-localization is unlikely to occur by chance (*P*=8.2×10^−10^, one-sided binomial test), indicating that NCOs are strongly enriched in CO hotspots, and providing further validation that the detected NCO events are real.

The enrichment of NCO in regions with high rates of meiotic CO suggests that the NCO resolutions are meiotic in origin. To explore this question further, we compared the locations of the NCO events with a recently reported genome-wide map of meiotic DSB hotspots in human spermatocytes [16]. We focused our analysis on NCO events transmitted by individuals likely to carry only the *PRDM9* zinc finger A or B alleles (see Methods “PRDM9 variants”), since individuals with different *PRDM9* zinc finger domains are known to have hotspots in distinct locations [23]. We further omitted NCO sites that occur near COs (and are consequently ambiguous as to which homolog converted; see below and Methods “Inclusion criteria”). For this analysis, we analyzed NCO events rather than single NCO sites, and report an event as overlapping a DSB if any of NCO site within it overlaps a DSB. By these criteria, there are 51 events, of which 26 (51%) overlap a meiotic DSB hotspot. Moreover, when focusing on events transmitted by males (because the DSB map is for spermatocytes), 19 of 27 events (70%) overlap a DSB hotspot. This enrichment is highly significant, as only 5.5% of informative sites overlap a DSB hotspot (P<10^−8^, calculated from 10^8^ permutations). Thus, the NCOs tend to occur at sites of meiotic DSB.

Moreover, the rate at which the NCO events overlap (sex-averaged) historical hotspots inferred from LD is almost identical to the rate at which meiotic DSBs occur in such locations. Considering all unambiguous NCO event locations in male *PRDM9* A/B-only carriers, 56% (15/27) overlap the (population-averaged) LD-based hotspots [24], when between 52% and 63% of DSB hotspots from spermatocytes do, depending on the source population of the LD map analyzed [16]. The overlap for NCO events from both sexes is similar, with 55% (28/51) of events overlapping LD-based hotspots. Finally, there is no overlap of the NCO events with putative fragile sites [25], the likely hotspots for mitotic recombination (see [26]). Given these observations, we conclude that most (possibly all) of our events arose in meiosis.

### Rate of non-crossover events and their location in the two sexes

The observation of 97 ascertained NCO sites out of 12.1 million informative sites provides an estimate of the rate of NCO per bp. Assuming the set of informative sites is unbiased with respect to the recombination rate, the rate of NCO is equivalent to the number of sites affected by NCO divided by the number of informative sites. This represents the proportion of the genome affected by NCO, or equivalently the probability that a given site will be part of a NCO tract per meiosis.

As Figure 2a shows, however, our SNP array data are enriched for regions of high recombination relative to the full genome, and it is necessary to account for this bias. We therefore estimated the rate of NCO in each of six recombination rate intervals based on the HapMap2 recombination map (Figure 2a), by dividing the number of NCO sites by the number of informative sites observed in each bin. The overall NCO rate is then the sum of these rates, each weighted by the proportion of the autosomes that occurs in the bin. This procedure yields a sex-averaged rate of *R*=5.7×10^−6^ per bp per meiosis (and a 95% confidence interval [CI] of 4.5×10^−6^ – 7.3×10^−6^, calculated by 40,000 bootstrap samples with 10 Mb blocks).

**Figure 2.**
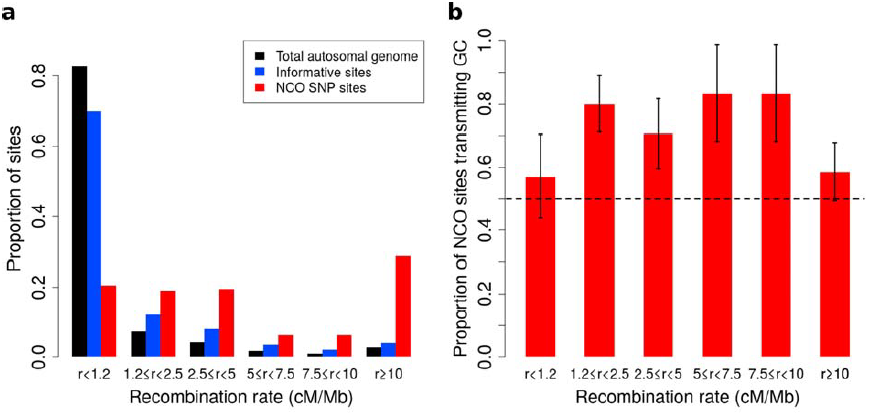
Proportion of non-crossover sites and rate of GC vs. AT allele transmissions across recombination rate bins. **a**, Histogram of proportions of sites that fall into six ranges of recombination rates [24] for the autosomal genome, all informative sites, and the identified NCO sites (see Methods “Crossover and recombination rates”). **b**, Rate of transmissions of G or C at AT/GC SNPs, across six recombination rate bins. Overall Plot shows standard error bars.

Sperm-typing data have also been used to examine the number and tract length of NCO events. Notably, a study by Jeffreys and May that examined three hotspots in detail [5] found the number of NCO events to be 4–15 times that of COs, with a mean tract length of 55–290 bp. The rate *R* can be calculated as the number of NCO tracts in a meiosis multiplied by the tract length and divided by the genome length. Using the estimates from Jeffreys and May yields *R*=2.6×10^−6^ to 5.2×10^−5^/bp/generation (for a genome-wide CO rate of 1.2 cM/Mb), a range that includes our estimate. Our results are therefore concordant with those from sperm-based analyses; they are also consistent with several LD-based studies of genome-wide levels of NCO [6,7,10].

Considering the parent of origin of each NCO event, we found that the two SNP arrays differ significantly in number of events detected per sex (*P*=5.1×10^−4^, *χ*^*2*^ 1 degree of freedom [df] test), with the lower density SNP dataset uncovering fewer male-specific events than expected. This bias may be caused by a lower coverage of the telomeres in the low density SNP array, and makes the analysis of potential differences in NCO rate between the sexes difficult. Nevertheless, considering the position of events captured by genotype arrays reveals broad-scale localization differences, with male events more prevalent in the telomeres and female events relatively dispersed throughout the genome (Figure 1c,d). These sex differences in localization are similar to those seen for CO events [12], as expected from a shared mechanism for the broader-scale (e.g., megabase-level) control of both types of recombination.

### GC-biased gene conversion

Deviations from the Mendelian expectation of 50% transmission of each allele at a polymorphic site have been observed a a number of recombination hotspots in humans. Many of these asymmetries result from polymorphisms that occur within motifs bound by PRDM9 [14]. Recombinations at these sites typically show under-transmission of the allele that better matches the *PRDM9* motif, a phenomenon thought to arise through initiation bias due to more frequent breakage of the homolog with a better match to the motif. We identified four NCO events that overlap sequences that match at least six of the eight predictive bps in the degenerate 13-mer motif bound by PRDM9 [27] (in all four cases, there are exactly 6 of 8 matching bps) and in which a SNP occurs at one of the non-degenerate positions. Because the *PRDM9* motif is GC rich, initiation bias would be expected to predominantly transmit AT alleles, but instead all four of these events transmit GC alleles. Notably however, for three of the events, sequences that match the *PRDM9* motif at 7 of 8 positions occur within 2 kb on the NCO site, and for the fourth, another motif with 6 of 8 bps matching occurs within 2 kb. Thus, these four events may not be caused by initiation bias.

A distinct form of bias in transmission that does not depend on the presence of polymorphisms in the *PRDM9* binding motif is thought to occur when AT/GC heteroduplex DNA arises during the resolution of recombination and is preferentially repaired towards GC alleles [15]. A recent sperm-typing study reported on two loci that exhibit such biased gene conversion, associated with NCO but not CO events [11]. This sperm-based study is, to our knowledge, the first to demonstrate direct evidence of gBGC in mammals. In the NCO events identified here, we saw no evidence for a difference in GC transmission rate between the two SNP density datasets (*P*= 0.18, *χ*^*2*^ 1-df test), or between males and females (*P*=0.79, *χ*^2^ 1-df test), and so considered the data jointly. For this calculation, we again omitted the ambiguous NCO events (described below and Methods “Inclusion criteria”) and we excluded the four sites that occur within *PRDM9* motifs. The remaining 92 NCO sites all have an AT allele on one homolog and GC on the other, a consequence of the fact that only ≤ 1% of sites on the Illumina SNP arrays are A/T or C/G SNPs. We observed a strong bias towards the transmission of G or C: Of the 92 sites, 63 transmit G or C alleles (68%, 95% CI 58–78%; *P*=5.1×10^−4^, two-sided binomial test). SNP variants at CpG dinucleotides account for 39 of these 92 sites, and these also show GC bias, with 25 CpG sites (64%) transmitting GC alleles, and no evidence of rate difference between transmissions at CpG and non-CpG sites (*P*=0.58, *χ*^2^ 1-df test). By comparison, the sperm-typing study noted above found that 2 of 6 assayed hotspots exhibited detectable levels of gBGC, and these two loci transmitted GC alleles in ~70% of NCO transmissions [11]. Across recombination rate bins, we observed consistent GC transmission rates (*P*=0.46, *χ*^2^ 5-df test Figure 2b). Since the strength of gBGC depends on both the degree of bias and the rate of recombination, this finding implies that the effects of gBGC will be strongest in high recombination rate regions, as seen in analyses of polymorphism and divergence [15].

### Non-crossover gene conversion tract lengths

The data allow us to estimate NCO tract lengths, with upper bounds derived from informative SNPs that flank a NCO tract and lower bounds given by the distance spanned by SNPs involved in the same tract. As previously noted, most NCO events involve only one SNP, but a total of twelve regions (ten with information from SNP array data only, and two including information from the sequence data) have tracts that include multiple SNPs (as plotted in Figure 3). From these data, we deduced that five of these events have a lower bound on tract length of at least 1 kb while the smallest is at least 94 bp. In turn, one tract is at most 144 bp—only slightly longer than the minimum tract involving more than one SNP (≥ 94 bp)—and four events have tracts that must be shorter than 1,400 bp. These observations, coupled with the variable length in tracts that occur in the clustered NCO events described below (see Figure 4a), suggest that tract lengths span at least an order of magnitude (i.e., 100-1000 bp).

**Figure 3.**
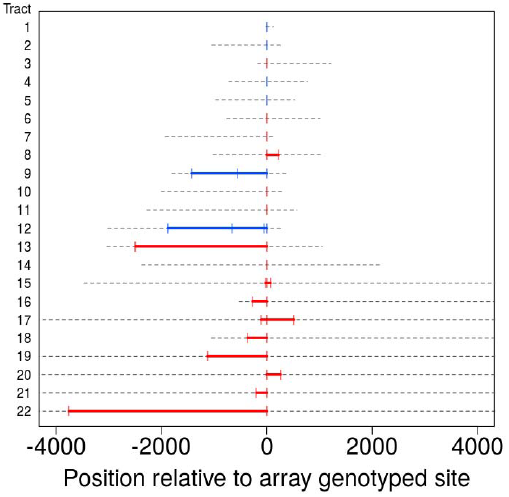
Tract lengths for identified non-crossovers. Tract lengths for the 22 NCO events that either have 2 or more SNPs in a tract or have maximum length of ≤ 5 kb. Each line corresponds to a NCO tract; lower bounds on length appear in color, with red corresponding to tract lengths informed by SNP array data and blue corresponding to tract lengths from sequence data. Gray dashed lines represent the region of uncertainty surrounding the tract length, with the end points being the upper bound on tract length. Tracts are sorted by the upper bound on tract length.

**Figure 4.**
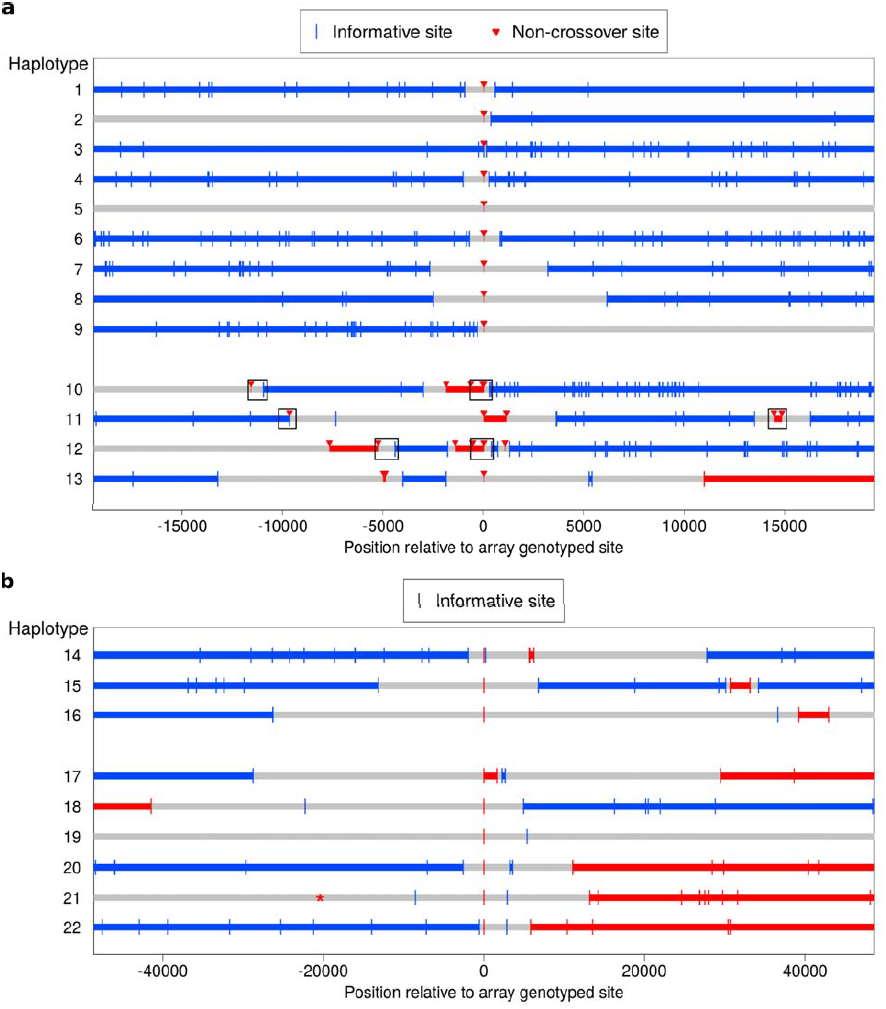
Clustered non-crossover events evident in resequencing data. **a**, Recombination patterns in whole genome sequence data for the region surrounding 13 NCO events originally identified in the SNP array data. Each horizontal line represents a haplotype transmission from a single meiosis, and position 0 on the x-axis corresponds to NCO sites identified in the SNP array data. Blue lines depict haplotype segments that derive from the parental homolog transmitted in the wider surrounding region, with blue vertical bars depicting informative sites. Red lines depict segments from the opposite homolog and are putative NCO events, with red arrows indicating informative sites. Grey lines are regions that have ambiguous haplotypic origin. For haplotypes 1–9, only a single site exhibits NCO. For haplotypes 10–13, several NCO sites appear in a short interval near each other but separated by informative SNPs from the background haplotype. Boxes indicate regions for which we preformed Sanger sequencing (see text). **b**, Clustered recombination events identified in the SNP array data; note the different scale on the x-axis compared with panel **a**. Here, haplotypes 14–16 are clustered NCO events while haplotypes 17–22 occur near but not contiguous with CO events (note the switch in haplotype color between the left and right side of the plot). It is uncertain whether the alleles descending from the blue or the red haplotype represent NCO events (Methods “Inclusion criteria”); thus the plot uses the same symbol for informative sites from both parental haplotypes. Haplotype 19 also appears to have resulted from a CO, but with informative sites more distant than the range of the plot. Haplotype 21 contains an informative marker that is ambiguous in the third generation and therefore was not detected initially, but it is plotted here with a * symbol. The ambiguous phase in the third generation is consistent with neighboring sites and not indicative of an incorrect genotype call.

Because NCOs identified using SNP arrays are sparsely sampled, our data may be enriched for events with longer tracts, because such tracts impact a larger number of sites. This effect would bias an estimate of the mean tract length using the data from this study. It is also possible that some of the longer events result from clustered but disjoint tracts, as described below.

### Complex clustered non-crossover tracts in sequence and SNP array data

We used Complete Genomics resequencing data generated by the T2D-GENES Consortium to examine variants surrounding several of the identified NCO events at closer resolution. In order to confidently phase these regions, we required sequence data for both parents and three children (including the NCO event recipient); such data were available for two pedigrees. In these pedigrees, there are a total of 15 regions with evidence for a NCO event in the SNP array data. Two of these regions are not included in this analysis: for one, the sequence data do not contain a genotype call for the site putatively affected by NCO, while in the other, genotype calls do not match the sequence data. Neither locus contains other sites affected by NCO in the sequence data.

Figure 4a shows the phase for the 13 regions included. In four cases (haplotypes 10–13), multiple disjoint NCO tracts occur within a short interval of less than 30 kb, with the discontinuities evident from informative sites located between the NCO tracts. Two of these events (haplotypes 11 and 13) occur near COs, and the sites that experienced NCO are ambiguous; Figure 4a plots the NCOs that result in shorter tracts. The four cases occur in a single pedigree, three in the mother, and one in the father (haplotype 11). The LD-based genetic map length of the 100 kb around these four regions ranges from 0.034 cM to 0.28 cM. Using these genetic lengths to estimate the probability of NCO initiation (Methods “Examination of regions containing clustered non-crossovers”), we found that this clustering is highly unexpected, with a probability of observing two independent tracts within the four 100 kb regions ranging from *P*=2.6×10^−6^ to 1.7×10^−4^ (for each region independently).

To check for possible artifacts, we performed Sanger sequencing of the three-generation pedigrees for six regions in three of these four haplotypes, indicated by boxes in Figure 4a. The Sanger sequence data are concordant with the genotypes from the whole genome sequence data at every site and in all individuals for which we were able to call genotypes (see Methods “Examination of regions containing clustered non-crossovers”). Moreover, we checked for overlap between these regions and the following resources: (a) recent segmental duplications that have divergence between them of <2% [28]; (b) the 35.4 Mb “decoy sequences” released by the 1000 Genomes Project [29] which contain regions of the genome that are paralogous to sequence from Genbank [30] and the HuRef alternate genome assembly [31]; and (c) regions of the genome with excess read mapping in the 1000 Genomes Project [32]. Our quality control procedure already removed individual SNPs that overlap several of these resources (Methods “Quality control procedures”), and this analysis showed no overlap for non-genotyped sites within the regions containing these clustered events. Finally, to evaluate whether an unknown paralogous sequence variant could confound the results, we considered the genotype status (homozygous or heterozygous) of variants within these regions. Haplotypes 10 and 12 include heterozygous and homozygous genotypes both within and outside the NCO tracts. For haplotypes 11 and 13, genotypes at all NCO sites are homozygous whereas the genotypes at other sites (blue in Figure 4a) are heterozygous. This observation raises the concern of a structural variant or duplicated sequence that spans the nearby CO breakpoint, leading heterozygous genotypes to be mismapped and possibly to mimic a NCO tract. Reassuringly, at all these positions, the non-transmitting parent is homozygous, one sibling is heterozygous, the other is homozygous, and neither of the other children received a recombinant haplotype. Thus, all four events appear to comprise true NCOs, but caution is warranted in interpreting two of the four cases.

Intriguingly, the alleles within the NCO events show strong GC bias: For the two unambiguous events (haplotypes 10 and 12), GC alleles were transmitted at 9 out of 10 heterozygous AT/GC SNPs affected by NCO. Moreover, considering all sites in haplotypes 10–13 irrespective of NCO status, G or C was transmitted at 29 of 37 (78%) heterozygous AT/GC SNPs. These findings raise the possibility that the patchy repair resolutions observed in the four events result from a GC biased repair process that operates discontinuously within long stretches of heteroduplex DNA. Alternatively, these results could be explained by repeated template switching, as has been observed in *S. cerevisiae* [33].

The close clustering of NCO events occurs in 4 of 15 (27%) cases that we were able to examine. As in the case of long tracts, however, our sparse, SNP array-based sampling may be more likely to detect clustered NCO events (since multiple tracts may affect a larger proportion of sites), and therefore the rate of clustering is likely to be lower than observed here.

Further examination of our array-based data revealed additional events: three more clustered NCO events as well as six NCO events near but disjoint from CO resolutions (Figure 4b). Two of these haplotypes (numbers 18 and 19) are the same cases that show clustered NCO in sequence data (Figure 4a, haplotypes 11 and 13); all other events were seen in distinct pedigrees. These complex CO resolutions shed light on the distances over which such events may occur. The complex CO events previously described in humans were seen in assays of relatively short intervals of ≤ 4 kb around CO breakpoints, and yielded an estimated frequency of 0.17% [14]. The results from the current study indicate that complex resolutions also occur farther from the CO breakpoint, so may be more common. Whether the observations at short and longer distances result from the same phenomenon remains to be elucidated.

To our knowledge, this is the first observation of clustered but discontinuous NCO gene conversion tracts in mammals, although patterns that resemble those shown in Figure 4a have been reported in meiosis [34,35] and mitosis [36,37] in *S. cerevisiae.* This phenomenon and the distant forms of complex CO both point to a property of mammalian recombination that is poorly understood and in need of further characterization.

### Contiguous and clustered recombination events spanning larger distances

In addition to the NCO events with tracts that span no more than 5 kb, we identified five longer-range recombination events: three continuous tracts, and two that showed a clustering pattern (see Figure 5). Each event occurred in a different pedigree; the continuous tract that spans ~79 kb was transmitted by a male, and the four other events occurred in females. The long continuous tracts could conceivably reflect double COs in extremely close proximity, as might arise from a CO-interference independent pathway [38], but the clustered events cannot be explained in this way. For two events, sequence data are available and validate the genotype calls, indicating that the case that spans at least 9 kb in the genotype data is in fact at least 18 kb long (haplotype 23), and confirming the case in which clustered events span ~203 kb (haplotype 26).

**Figure 5.**
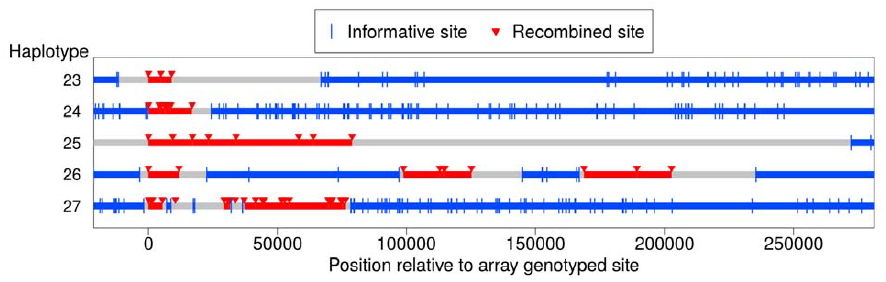
Long-range recombination events observed in sequence data. Shown are three contiguous recombination tracts with length ≥ 9 kb, ≥ 16.9 kb, and ≥ 79 kb as well as two sets of clustered long-range recombination events that span ~200 kb and ~76 kb.

Haplotypes 23, 24, and 27 reside on the p arm of chromosome 8 where a long inversion polymorphism occurs [39]. Single COs within inversion heterozygotes can be misinterpreted as more than one CO event [40], yet these three recombination events are > 1.7 Mb outside the inversion breakpoints, so should not be affected. One possibility is that the large inversion polymorphism leads to aberrant synapsis during meiosis, leading to complex repair of DSBs. In that regard, we note the transmitter of haplotype 23 is heterozygous for tag SNPs for the 8p23 inversion polymorphism [39], and that a sibling inherited a haplotype from the same parent with a CO at the same position as the end of the tract for haplotype 23. This co-localization may be due to effects of the inversion on synapsis; alternatively, this could indicate that the sites are incorrectly positioned, resulting in inaccurate inference of breakpoint locations [40]. The pattern is haplotype 27 is even more complex and difficult to explain.

## Discussion

NCO gene conversion reshuffles haplotypes and shapes LD patterns, at a rate that we estimate to be 5.7×10^−6^/bp/generation. This suggests that roughly 16,530 (95% CI 13,050–21,170) sites will be affected by NCOs in each generation (for a euchromatic genome length of 2.9×10^9^ bp). If the average tract length were 75 bp (consistent with [4,5]), ~220 NCO events (95% CI 174–282) are expected to occur in each generation. Given that the sex-averaged number of COs is ~30 each generation (e.g., [41]), the number of NCOs that we detect is thus in rough agreement with a 10:1 NCO to CO ratio genome-wide [8,9].

NCO events only impact variation patterns when they occur at heterozygous sites, so in many contexts, this rate is most of interest when scaled by human heterozygosity levels (i.e., the proportion of sites that differ between two homologous chromosomes). Assuming that the heterozygosity rate is π = 10^−3^ [42], roughly 17 (95% CI 13–21) variable sites are expected to experience NCO in each meiosis. This estimate is on the same order as the number of sites affected by *de novo* mutation in each generation [43].

In regions that experience NCO, our results indicate that there is frequent over-transmission of G or C alleles. Indeed, we observed GC transmission in 68% of events (95% CI 59–78%), with no difference in the rate of gBGC across a range of recombination rates (Figure 2b). More generally, our results provide a direct confirmation of the presence of gBGC, and lend strong support to the hypothesis that it could play a major role in shaping base composition over evolutionary timescales [15]. Our estimated rate of GC transmission is high relative to what was found in the recent sperm-typing study of a handful of hotspots [5]. In that regard, one possible caveat is that, under certain conditions on mutation, the ascertainment bias of SNP genotyping arrays could lead SNPs subject to stronger biased gene conversion to be enriched, and thus lead us to slightly over-estimate the strength of biased gene conversion across the genome.

Interestingly, a recent reanalysis of data from *S. cerevisiae* [44] showed that in this species of yeast, gBGC is associated with CO gene conversions but not NCOs, with a GC transmission rate of ≤ 55% [45]. Both our findings and recent results from human sperm [11] indicate that in contrast, in humans, gBGC does operate in NCO events, pointing to a difference in repair mechanisms between humans and yeast that remains to be elucidated.

Considering the distribution of SNPs in NCO tracts, we found lengths that vary over more than an order of magnitude, from hundreds to thousands of base pairs. Intriguingly, we also identified several examples of loci where multiple NCO tracts cluster within 20–30 kb intervals, as well as instances of complex CO over extended intervals. As a potential example of the same phenomenon, a study of *de novo* mutations reported observing regions with NCO sites across intervals spanning between 2–11 kb [46]. These events may either be long NCO tracts or clustered but discontinuous NCO events in the same meiosis. In any case, the complex NCO resolutions seen in our pedigree data has not been reported in mammals previously, and is consistent with either with patchy GC biased repair across long stretches of heteroduplex DNA or repeated template switching during the repair of DSBs. Understanding their source will be important for studies of mammalian recombination and for improving population genetic models of haplotypes and LD.

Going forward, whole genome sequencing of human pedigrees will enable unbiased analyses of *de novo* NCO at relatively high resolution. Of particular interest will be the estimation of the strength of gBGC free of ascertainment bias, as well as systematic examination of tract length distribution and the patterns of complex NCO resolutions revealed by this study.

## Methods

### Samples and sample selection

This study analyzed Mexican American samples from the San Antonio Family Studies (SAFS) pedigrees. SNP array data were generated for these individuals as previously described [19-21]. Our study design required the use of three-generation pedigrees with SNP array data for both parents in the first generation, three or more children in the second generation, one or more grandchildren, and data for both parents for any included grandchildren. Within the entire SAFS dataset of 2,490 individuals, there are 35 three-generation pedigrees consisting of 496 individuals that fit the requirements of this design. As noted below, one of these pedigrees was not included in the analysis, so the overall sample consists of 34 pedigrees and 482 individuals.

Each sample was genotyped using one of the following Illumina arrays: the Human660W, Human1M, Human1M-Duo, or both the HumanHap500 and the HumanExon510S (these latter two arrays together give roughly the same content as the Human1M and Human1M-Duo).

Most of the samples—21 out of the 34 analyzed pedigrees containing 293 individuals—have SNP data derived from arrays with roughly equivalent content and ~1 million genotyped sites. We analyzed all these samples across the SNPs shared among these arrays, with data quality control applied collectively to all samples and sites (see below). After quality control filtering, 896,375 autosomal SNPs remained for the analysis of NCO.

Data for the other 13 out of 34 analyzed pedigrees comprise 189 individuals and were analyzed on a lower density SNP arrays. The majority of the samples in these pedigrees (105 individuals) have SNP array data from ~660,000 genotyped sites. The other samples (84 individuals) have higher density genotype data available, but because other pedigree members have only lower density data, we omit these additional sites from analysis. After quality filtering, this lower SNP density dataset contained 513,283 autosomal sites.

### Quality control procedures applied to full dataset

Initially, sites with non-Mendelian errors, as detected within the entire SAFS pedigree, were set to missing. We next ensured that the locations of the SNPs were correct by aligning SNP probe sequences to the human genome reference (GRCh37) using BWA v0.7.5a-r405 [47]. Manifest files for each SNP array list the probe sequences contained on the array and we confirmed that these probe sequences are identical across all arrays for the SNPs shared in common among them. We retained only sites that (a) align to the reference genome with no mismatches at exactly one genomic position and that (b) do not align to any other location with either zero or one mismatches.

We updated the physical positions of the SNPs in accordance with the locations reported by our alignment procedure and utilized SNP rs ids contained in dbSNP at those locations. We omitted sites for which multiple probes aligned to the same location. Some sites had either more than two variants or had non-simple alleles (i.e., not A/C/G/T) reported by dbSNP, and we removed these sites. We also filtered three sites that had differing alleles reported in the raw genotype data as compared to those reported for the corresponding sites in the manifest files. We filtered a small number of sites for which the manifest file listed SNP alleles that differed from those in dbSNP at the aligned location.

Some SNPs are listed in dbSNP as having multiple locations or as “suspected,” and we removed these sites from our dataset. We also removed sites that occur outside the “accessible genome” as reported by the 1,000 Genomes Project [42] (roughly 6% of the genome is outside this), and sites that occur in regions that are segmentally duplicated with a Jukes-Cantor K-value of <2% (this value closely approximates divergence between the paralogs) [28]. Finally, we removed sites that occur within a total of 17 Mb of the genome that receive excess read alignment in 1,000 Genome Project data [32].

We next conducted more standard quality control measures by performing analyses on two distinct datasets: (1) including all individuals that were genotyped at ~1 million SNPs (1,932 samples) and (2) including all 2,490 samples. On the densely typed dataset, we first removed any site with ≥ 1% missing data and those for which a *χ*^2^ test for differences between male and female allele frequencies showed |Z|≥3. We then removed 29 samples with ≥2% missing data. Next we examined the principal components analysis (PCA) plots [48] generated using (a) the genotype data and (b) indicators of missing data at a site. These plots generally show an absence of outlier samples, and the genotype-based PCA plot appears consistent with the admixed history of the Mexican Americans (results not shown).

For the datasets that include samples typed at lower density, we first removed sites with ≥1% missing data and sites with male-female allele frequency differences with |Z|≥3. This filtering step yields SNPs of high quality that are shared across all SNP arrays, including the lower density Human660W array. Next we removed 30 samples with ≥2% missing data. Lastly, we examined PCA plots generated using (a) genotype and (b) missing data at each site, and these plots are again generally as expected with an absence of outlier samples (results not shown).

### Phasing and identifying relevant recombination events in three-generation pedigrees

We performed minimum-recombinant phasing on the three-generation pedigrees using the software HAPI [22], but with minor modifications because this program phases nuclear families independently. Specifically, our approach phased nuclear families starting at the first generation family. After this completed, we phased the families from later generations while utilizing the haplotype assignments from the first generation. Our approach assigned the phase at the first heterozygous marker to be consistent across generations in the individuals shared between the two nuclear families. (Shared individuals are members of the second generation who are a child in one family and a parent in another.) This approach helps produce consistent phasing across generations and does not introduce extra recombinations since the phase assignment at the first marker on a chromosome is arbitrary.

After phasing, our method for detecting NCO events also handled sites with inconsistent phase between the families (though in practice nearly all sites have consistent phase assignments between families). This method excluded sites that have inconsistent phase and that occur within a background of flanking markers with consistent phase; we examined these sites individually and confirmed that they do not represent NCO events, but are likely driven by genotyping errors. When 10 or more informative SNPs in succession are inconsistent across families, we assumed that a CO event went undetected in one of the generations, and inverted the phase for the relevant individuals in order to identify putative NCO events.

We analyzed the inferred haplotype transmissions to identify sites that exhibit recombination from one haplotype to the other and then back again. The detection approach identified any recombination events that switch and revert back to the original haplotype within ≤20 informative SNPs.

### Pedigree-specific quality control and determination of informative sites

Genotypes are only informative for which haplotype a parent transmits—and therefore recombination—at sites where the parent is heterozygous. We employed a pedigree-specific quality control measure by only considering sites in which all individuals in the full three-generation pedigree have genotype calls and no missing data; other sites are omitted. This requirement helps address possible structural or other complex variants that are specific to a particular pedigree and that may adversely affect genotype calling (as evidenced by a lack of a genotype call for some individual in that pedigree at the given site).

Because NCOs occur relatively infrequently, it is unlikely that the same position will experience NCO in multiple generations. We therefore excluded sites that exhibit NCO in any grandchild (i.e., locations with potential NCO events transmitted from the second generation). We applied this filter regardless of the NCO status in earlier generations in order to obtain unbiased ascertainment of events and informative sites. We also excluded sites that exhibit potential NCO events from a given parent and where that parent only transmits one haplotype. In this case, the genotype from the transmitting parent is likely to be in error and to be homozygous; given this consideration, we considered the site as invalid for both parents.

In principle, all children in the second generation are useful for studying meiosis in their parents, but to reduce false positives, we only analyzed a subset of these children. Specifically, we only analyzed a child if data for his/her partner and one or more of their children (grandchildren in the larger pedigree) were available.

We counted a site as informative (or not) relative to a given parent and a given child if sufficient data for relatives were available and if it satisfied six requirements. First, we required the parent to be heterozygous at the site. Second, as shown in Figure 1b, we required the allele that the given parent transmitted to the child also be transmitted to at least one grandchild. Third, in any series of otherwise informative sites, we counted all but the first and last sites as informative since we detect NCO events as haplotype switches relative to some previous informative site. Fourth, except at sites that are putatively affected by NCO, we required that a second child to have received the same haplotype as the child that is potentially informative. This requirement helps to ensure the validity of the heterozygous genotype call of the parent. As an example, consider a pedigree with four children, three of whom received a haplotype ‘A’ at some site and the fourth of whom received haplotype ‘B’. If the fourth child were to receive a NCO at some subsequent position, it would receive haplotype ‘A’, and thus all four children would receive the same haplotype. This scenario violates the requirement that the alternate allele (from the haplotype not affected by NCO) be transmitted to at least one second-generation child. Thus, in this example, the fourth child is not informative at this example site (where it is the sole recipient of haplotype ‘B’). Note however that this site could be informative in the other children if they meet the other requirements listed here.

Fifth, we required that the site be phased unambiguously across two generations, and that if a NCO had occurred, the phase at the site would remain unambiguous in the first generation. Sites in which all individuals in a nuclear family are heterozygous have ambiguous phase. Thus, if a given child is homozygous at a marker but all other individuals in the family are heterozygous, the child is not informative at that site since a NCO event would lead the child to be heterozygous. We note that it is possible to identify putative NCOs when a child receives a haplotype that has recombined from otherwise ambiguous phase to be homozygous at this type of marker. Indeed, we identified five such putative NCO sites, but did not include them when calculating the rate of NCO since the denominator does not include ambiguously phased sites and is therefore ascertained differently.

Finally, we imposed further conditions on the transmitted haplotypes and the genotype calls in the third generation. Our focus for these filters was the case in which a NCO recipient is called homozygous but is truly heterozygous, and his/her partner and children are all heterozygous. In this case, the phasing procedure may incorrectly infer that an allele transmitted by the recipient was instead transmitted by his/her partner; thus the potential NCO allele is not necessarily observed in the grandchildren. To address this issue, we filtered sites that have two properties: (1) the recipient is called homozygous, but the partner and grandchildren are all heterozygous, and (2) both parents transmit only one of their haplotypes to the third generation. If, in contrast to property (2), either the recipient or the partner transmits both his/her haplotypes, this type of erroneous NCO genotype call will produce haplotype assignments in the grandchildren with apparent recombination events relative to flanking markers. As a result, there will be an apparent NCO in the third generation and a filter noted above will remove the site from consideration.

### Pedigrees included in the analysis

We excluded one of the 35 available three-generation pedigrees from our analysis. The NCO recipient in this pedigree has a missing data rate that is more than double any other NCO recipient, suggesting genotype quality issues; accordingly, we observed an excessive rate of NCO event calling in this pedigree (results not shown).

### Quality filtering of double recombination events in close proximity

Our method identified all double recombination events (defined as switches from one haplotype to the other and then back again) that span 20 informative sites or fewer. We examined the haplotype transmissions at each such reported event by hand to ensure that segregation to all children and grandchildren matches expectations. A few sites exhibited NCO events in the same interval in two or more children. Because NCO is relatively rare, it is unlikely that these are true events. Additionally, some sites were consistent with NCO events transmitted to the same child from both parents; these are again unlikely to be real and are more likely caused when a child is homozygous for one allele but called homozygous for the opposite allele. We therefore considered these cases false positives.

Although we omitted sites in which grandchildren exhibit putative NCO events that occur at a single site, the software did not filter putative NCOs that span multiple sites. We examined all events by hand, and excluded three reported NCO events in which the grandchildren either exhibit putative NCOs longer than one SNP (therefore undetected) or show aberrant genotype calls.

The main text describes five long-range recombination events shown in Figure 5. For all these events, the recombined alleles at every site were transmitted to the third generation with no apparent recombinations or NCO events in the third generation. We excluded two other events with unexpected transmissions to the grandchildren. Specifically, one 4-SNP contiguous tract shows transmission to the third generation for three of the four recombined SNPs, but one SNP in middle of the tract was not transmitted and shows an apparent NCO in the third generation. The other 18-SNP long contiguous tract shows a putative NCO transmitted from the opposite parent across this same interval. We also excluded an event in which two sites separated by ~27 kb exhibit NCO in the second generation, but where one site has ambiguous phase in the third generation and would not be expected to have such phasing on the basis of flanking markers.

### Validating non-crossover events

We tested for overrepresentation of either heterozygous or homozygous genotype calls in the recipient of the putative NCOs. Overrepresentation would suggest bias and possibly artifactual detection of NCOs, but we saw no evidence of bias (*P*=0.56, two-sided binomial test). This analysis excludes the five sites identified using non-standard ascertainment and which are homozygous by detection, and also excludes a sixth non-standard site (described below in “Inclusion criteria”).

Of the 482 individuals that we analyzed using SNP array data, 98 were whole genome sequenced by the T2D-GENES Consortium and we were therefore able to check concordance of genotype calls. We attempted validation on all sites for which data were available for the transmitting parent or a recipient (either the child or a grandchild) of the putative NCO site (Supplementary file 1). Within these 98 samples, genotype calls were available for 52 of the putative NCO sites (of the 103 total); 42 of these sites include data for both the transmitting parent and a NCO recipient. One additional site had data available for relevant samples, but the sequence data do not contain calls for that position. We compared genotypes for every available parent, child, partner of the NCO recipient, and children of the recipient (grandchildren in the larger pedigree). For ambiguous NCOs, we required data for both possible NCO orientations to be concordant in order to count as validated. The genotype calls for all inspected individuals are concordant between the two sources of data for 50 of the 52 sites. One of the inconsistent sites shows a discordant genotype call between the datasets for the recipient of the NCO, but a concordant call for his child (the grandchild in the pedigree). This inconsistency suggests that the genotype data may in fact be correct. The other discrepancy occurs at a site where sequence data were unavailable for the recipient of the NCO. Here, the genotype call for the transmitting parent is discordant between the two sources of data, and the error source is ambiguous; we retained this site in the analyses.

### Crossover and recombination rates

CO rates are those reported by deCODE [12] based on COs detected in large Icelandic pedigrees. The original map is reported for human genome build 36 and was lifted over to build 37 coordinates. This map is estimated to have resolution to roughly 10 kb, and we therefore computed recombination rates in cM/Mb at each site using the genetic distances from the map at the 10 kb surrounding a site and divided by this (10 kb) window size. Because this map omits relatively large telomeric segments, we did not have rates for many sites from the SNP arrays and from the identified NCO events. We used linear interpolation to obtain rates at sites within the range of the map but not directly reported. The proportion of sites in the “autosomal genome” in Figure S1 derives from all sites within the reported positions in the autosomal genetic map.

The HapMap2 LD-based recombination rates are from the genetic map generated by the HapMap Consortium [24] using LDhat [49] that was subsequently lifted over to human genome reference GRCh37. We used analogous methods for calculating recombination rates from this map as for the CO map mentioned above, including a window size of 10 kb and linear interpolation. A few sites on the higher density SNP data (12 of 896,387) fall outside the interval of positions reported in the map and were not included in our analyses.

### PRDM9 variants in the sample

Mexican Americans were previously shown to carry primarily *PRDM9* A and B alleles [50], and admixture with African descent groups may have led to the presence of *PRDM9* C variants. The derived allele at SNP rs6889665 is in strong LD with this *PRDM9* C variant: 96% of haplotypes with the ancestral allele contain < 14 zinc fingers, with most being A or B alleles; 93% of haplotypes with the derived allele contain *PRDM9* variants with ≥ 14 zinc fingers, including primarily the *PRDM9* C variant. With a larger number of zinc fingers, the PRDM9 C variant binds a degenerate 17 bp motif distinct from the motif bound by PRMD9 A and B [51]. The higher SNP density arrays include genotypes for this site, providing information about the likely *PRDM9* variant of the transmitting parent for 76/103 of the NCO sites. Of these, 11 events are transmitted by a likely *PRDM9* C carrier, with a total of five carrier parents within the 48 parents for which we have genotypes. The remaining 65 events are transmitted by individuals that are homozygous for the ancestral allele at rs6889665 and thus likely to carry only the PRDM9 A or B alleles, both of which bind the common 13-mer motif [23].

### Inclusion criteria for non-crossover and GC-bias rate calculations, hotspots, and tract lengths

Five NCO events were identified with a non-standard ascertainment and are inappropriate for inclusion in estimating the rate of NCO. A sixth, non-standard event is part of a three SNP long tract but has ambiguous phase in the third generation; it appears to be to be a NCO site on the basis of its presence in a tract and the fact that the ambiguous phase in the third generation is consistent with neighboring sites and not suggestive of artifact. None of these sites are expected to show bias with respect to allelic composition and we therefore included them when calculating the strength of GC-bias.

Somewhat more complex cases are NCO sites that occur near CO events (Figure 4b, haplotypes 17–22). In most, a single site appears to have been involved in the NCO event, and is followed by a single site that reverts to the first haplotype, and then by a CO. Depending on whether one considers the “background haplotype” to be the one upstream of the NCO and CO or downstream, the site in the NCO tract differs. Thus the sites affected by NCO are ambiguous. To simplify the examination of GC-bias, we excluded these sites from consideration. The excluded haplotypes are 17–22 (Figure 4b); haplotypes 11 and 13 are the same as 18 and 19 and are thus excluded, whereas haplotypes 10 and 12 and 14–16 are unambiguous NCO tracts and are included. (Haplotypes 23–26 in Figure 5 are long-range events that are not included in any analysis.) Additionally, to avoid confounding biased repair with initiation bias driven PRMD9 binding, we omit four events that overlap partial matches to the *PRDM9* motif as described in Results (“GC-biased gene conversion”).

To estimate the rate of NCO genome-wide, rather than exclude the ambiguous sites noted above—which would bias our rate calculation downwards—we instead included both possibilities in the rate calculation, and gave each of them a weight of 0.5, while other sites have a weight of 1. There are two effects of this weighting. First, if the recombination rate bin differs across these sites, they each contribute the weight of half a site to the rate calculation for those bins. Most sites fall into the same rate bin and therefore have the same effect as counting a single site. The second effect of weighting these sites is that, in one case, we cannot tell whether the NCO was 2 SNPs or only 1 SNP long. In this case, we counted the event as 1.5 NCO sites. Finally, we observed one instance of two adjacent putative NCO sites separated from a CO by three informative sites. The three informative sites span 19.6 kb—longer than our threshold for NCO events. In this case, we considered the two sites (which form a tract of length at least 264 bp) as part of a definitive NCO with weight 1.

For estimating the number of sites with CO rate ≥10 cM/Mb, we included only 1 SNP per tract and weighted ambiguous cases by 0.5 as above. Additionally, two ambiguous sites have CO rates that straddle this threshold, with one site slightly less, the other slightly more. To be conservative in estimating a P-value, we considered these sites as falling below the threshold.

We checked overlap between DSB hotspots determined by Pratto et al. [16] and the set of NCO sites that are unambiguous (i.e., omitting haplotypes 17–22 from Figure 4b) and for which the transmitting parent is likely to be homozygous for *PRDM9* A or B (see “PRDM9 variants” above). A secondary analysis of these DSB hotspots included only events transmitted by males. To calculate overlap with LD-based hotspots [24], we again included only unambiguous events but did not further restrict the analysis. To assess the significance of the overlap, we performed permutation by randomly sampling NCO sites among all informative sites, selecting the same number of adjacent SNPs as observed for each event. We repeated this process 10^8^ times, each time calculating the proportion of NCO events that overlapped a DSB. Out of 10^8^ permutations, no samples obtained at least the level of overlap seen for the actual NCO events; thus *P* < 10^−8^.

To examine tract lengths, we omitted all but one ambiguous event. For the one included ambiguous event, the two possibilities have tract lengths ≥1,615 bp and ≥365 bp (upper bounds are more than 25 kb for both). We included the shorter of these lengths (365 bp) since this lower bound holds for both possibilities. We note that the addition of the sixth, non-standard NCO site that is part of a three-SNP tract (see above) leads to a minimum tract length of 629 bp instead of 520 bp (obtained for the two-SNP tract identified with standard ascertainment).

### Examination of regions containing clustered non-crossovers

We calculated the probability of two NCO events occurring within the four intervals in which we observed clustered NCO by rescaling the genetic distances of these regions as reported in the LD-based map. (Note that this map includes some of the historical effects of NCO [52].) We earlier estimated the per bp rate of NCO *R*, and *R*=*N*×*l*/*G* where *N* is the number of NCO events that occur in a meiosis, *l* is the average tract length of these events, and *G* is the total genome length. The genome-wide average rate of *initiation* of NCO at a bp is simply *N*/*G* = *R*/*l*. For an interval with genetic map length *d* cM, we estimated the rate of initiating a NCO as *r*=*d*/*c*×*R*/*l*, where c=1.2 cM/Mb is the average genome-wide rate of CO, and where we assume *l*=75 bp. The probability of two independent NCO tracts (conservatively assuming lack of interference among events) is then *P*=*r*^2^. This calculation assumes the HapMap2 map accurately represents the relative rate of both CO and NCO events in an interval; a test for difference between the observed locations of NCO sites and expected locations based on this map are generally consistent with this assumption (*P*=0.23, *χ*^2^ 5-df test).

We performed Sanger sequencing on individuals from the three-generation pedigrees in which clustered NCOs occurred. Assayed samples included both parents, all children (including the NCO recipient), the partner of the NCO recipient, and all (four or five) grandchildren of that couple. Overall, sequencing included 11 or 12 samples for each of the three regions examined. We manually examined chromatograms to determine genotype calls. For haplotype 10 (Figure 4a), Sanger sequence data overlaps five SNPs called in the Complete Genomics data. For three of these five positions, the sequence data quality was sufficient to easily call genotypes in all samples, whereas for two positions, we called genotypes only in the four grandchildren. Three of the four grandchildren received the haplotype that resulted from a NCO, providing validation of the event at these sites. In all cases, the Sanger-based genotypes are concordant with the Complete Genomics genotypes. For haplotype 11, the Sanger sequence overlapped four SNPs called in the Complete Genomics data. For one of these sites, we called genotypes in all samples, and for two others, we omitted genotypes for one sibling of the recipient but called all other samples. For the fourth site, we could not determine the genotype of the transmitting parent and were uncertain of three of the four siblings of the recipient, but still obtained genotypes for the recipient, one sibling, and four grandchildren. At all four sites, the genotypes that we obtained are consistent with those in the Complete Genomics data. Finally, for haplotype 12, the Sanger sequence overlaps eight sites from the Complete Genomics data. We called genotypes in the five grandchildren at seven of the eight sites, and call four of the five grandchildren at the eighth site. Data quality for other individuals in the pedigree was high for four of the eight sites, but low for the other four sites. In all cases for which we obtained genotype calls, the Sanger data are concordant with the Complete Genomics data. Overall, the Sanger sequence data provided genotype calls for three (haplotypes 11 and 12) or four (haplotype 10) NCO sites (Figure 4a) as well as one (haplotype 11) and at least two sites (haplotypes 10 and 12) that descend from the background haplotype (red in Figure 4a).

The main text describes an additional analysis that checked the regions for potential mismapping from paralogous sequences elsewhere in the genome.

### Sanger Sequencing

We ran Primer3 (http://bioinfo.ut.ee/primer3/) using the initial presets on the human reference sequence from targeted regions to obtain primer sequences. For the suggested primer designs, we performed a BLAST against the human reference to ensure that each primer is unique, and ordered primers from Eurofins Operon. We tested each primer using the temperature suggested during primer design on DNA at a concentration of 10ng/uL and checked on a 2% agarose gel. For any primer with poor performance, we conducted a temperature gradient, and, if needed, a salt gradient until we found a PCR mix that performed well. Next we performed PCR on the samples of interest, running a small quantity on a 2% agarose gel. We then cleaned the PCR sample using Affymetrix ExoSAP-IT and ran sequencing reactions twice for each sample using Life Technologies BigDye Terminator v3.1 Cycle Sequencing Kit. Finally, we purified each sample using Life Technologies BigDye XTerminator Purification Kit and placed these onto the 3730xl DNA Analyzer for sequencing.

## Acknowledgements

We thank Scott Keeney, Maria Jasin, John Schimenti, Laure Ségurel, and Lorraine Symington for helpful discussions and Melanie Carless for bioinformatics support. We thank Swapan Mallick for sharing a version of the deCODE crossover map in GRCh37 coordinates. A.L.W. was supported by the NIH Ruth L. Kirschstein National Research Service Award number F32 HG005944 and by NIH GM83098 to M.P.. This work was partly completed while M.P. was a Howard Hughes Medical Institute Early Career Scientist. D.R. is a Howard Hughes Medical Institute Investigator. T2D-GENES project data generation was supported by NIH grants U01 DK085501, U01 DK085524, U01 DK085526, U01 DK085545, and U01 DK085584.

## Supplementary Material

**Figure S1.**
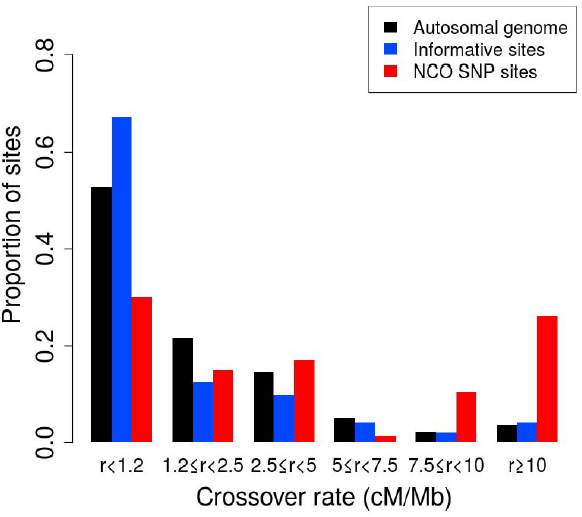
Proportion of non-crossover across crossover rate bins. Histogram of proportions of sites that fall into six ranges of crossover rates [12] for the autosomal genome, all informative sites, and the identified NCO sites (see Methods “Crossover and recombination rates”).

**Supplementary file 1. Non-crossover event details.** TSV file containing information about each NCO site. Descriptions of each column are listed as comments at the beginning of the file.

## References

1. Baudat F, Imai Y, de Massy B (2013) Meiotic recombination in mammals: localization and regulation. Nat Rev Genet 14: 794–806.

2. Cole F, Keeney S, Jasin M (2012) Preaching about the converted: how meiotic gene conversion influences genomic diversity. Ann N Y Acad Sci 1267: 95–102.

3. Youds JL, Boulton SJ (2011) The choice in meiosis – defining the factors that influence crossover or non-crossover formation. Journal of Cell Science 124: 501–513.

4. Cole F, Baudat F, Grey C, Keeney S, de Massy B, et al. (2014) Mouse tetrad analysis provides insights into recombination mechanisms and hotspot evolutionary dynamics. Nat Genet 46: 1072–1080.

5. Jeffreys AJ, May CA (2004) Intense and highly localized gene conversion activity in human meiotic crossover hot spots. Nat Genet 36: 151–156.

6. Ardlie K, Liu-Cordero SN, Eberle MA, Daly M, Barrett J, et al. (2001) Lower-Than-Expected Linkage Disequilibrium between Tightly Linked Markers in Humans Suggests a Role for Gene Conversion. Am J Hum Genet 69: 582–589.

7. Frisse L, Hudson RR, Bartoszewicz A, Wall JD, Donfack J, et al. (2001) Gene Conversion and Different Population Histories May Explain the Contrast between Polymorphism and Linkage Disequilibrium Levels. Am J Hum Genet 69: 831–843.

8. Baudat F, de Massy B (2007) Regulating double-stranded DNA break repair towards crossover or non-crossover during mammalian meiosis. Chromosome Research 15: 565–577.

9. Cole F, Kauppi L, Lange J, Roig I, Wang R, et al. (2012) Homeostatic control of recombination is implemented progressively in mouse meiosis. Nat Cell Biol 14: 424–430.

10. Gay J, Myers S, McVean G (2007) Estimating Meiotic Gene Conversion Rates From Population Genetic Data. Genetics 177: 881–894.

11. Odenthal-Hesse L, Berg IL, Veselis A, Jeffreys AJ, May CA (2014) Transmission Distortion Affecting Human Noncrossover but Not Crossover Recombination: A Hidden Source of Meiotic Drive. PLoS Genet 10: e1004106.

12. Kong A, Thorleifsson G, Gudbjartsson DF, Masson G, Sigurdsson A, et al. (2010) Fine-scale recombination rate differences between sexes, populations and individuals. Nature 467: 1099–1103.

13. Berg IL, Neumann R, Sarbajna S, Odenthal-Hesse L, Butler NJ, et al. (2011) Variants of the protein PRDM9 differentially regulate a set of human meiotic recombination hotspots highly active in African populations. Proceedings of the National Academy of Sciences 108: 12378–12383.

14. Webb AJ, Berg IL, Jeffreys A (2008) Sperm cross-over activity in regions of the human genome showing extreme breakdown of marker association. Proceedings of the National Academy of Sciences 105: 10471–10476.

15. Duret L, Galtier N (2009) Biased Gene Conversion and the Evolution of Mammalian Genomic Landscapes. Annual Review of Genomics and Human Genetics 10: 285–311.

16. Pratto F, Brick K, Khil P, Smagulova F, Petukhova GV, et al. (2014) Recombination initiation maps of individual human genomes. Science 346.

17. Auton A, Fledel-Alon A, Pfeifer S, Venn O, Ségurel L, et al. (2012) A Fine-Scale Chimpanzee Genetic Map from Population Sequencing. Science 336: 193–198.

18. Galtier N, Duret L (2007) Adaptation or biased gene conversion? Extending the null hypothesis of molecular evolution. Trends in Genetics 23: 273–277.

19. Mitchell BD, Kammerer CM, Blangero J, Mahaney MC, Rainwater DL, et al. (1996) Genetic and Environmental Contributions to Cardiovascular Risk Factors in Mexican Americans: The San Antonio Family Heart Study. Circulation 94: 2159–2170.

20. Duggirala R, Blangero J, Almasy L, Dyer TD, Williams KL, et al. (1999) Linkage of Type 2 Diabetes Mellitus and of Age at Onset to a Genetic Location on Chromosome 10q in Mexican Americans. The American Journal of Human Genetics 64: 1127–1140.

21. Hunt KJ, Lehman DM, Arya R, Fowler S, Leach RJ, et al. (2005) Genome-Wide Linkage Analyses of Type 2 Diabetes in Mexican Americans: The San Antonio Family Diabetes/Gallbladder Study. Diabetes 54: 2655–2662.

22. Williams A, Housman D, Rinard M, Gifford D (2010) Rapid haplotype inference for nuclear families. Genome Biology 11: R108.

23. Baudat F, Buard J, Grey C, Fledel-Alon A, Ober C, et al. (2010) PRDM9 Is a Major Determinant of Meiotic Recombination Hotspots in Humans and Mice. Science 327: 836–840.

24. The International HapMap Consortium (2007) A second generation human haplotype map of over 3.1 million SNPs. Nature 449: 851–861.

25. Fungtammasan A, Walsh E, Chiaromonte F, Eckert KA, Makova KD (2012) A genome-wide analysis of common fragile sites: What features determine chromosomal instability in the human genome? Genome Res 22: 993–1005.

26. Song W, Dominska M, Greenwell PW, Petes TD (2014) Genome-wide high-resolution mapping of chromosome fragile sites in Saccharomyces cerevisiae. Proceedings of the National Academy of Sciences 111: E2210–E2218.

27. Myers S, Freeman C, Auton A, Donnelly P, McVean G (2008) A common sequence motif associated with recombination hot spots and genome instability in humans. Nat Genet 40: 1124–1129.

28. Bailey JA, Gu Z, Clark RA, Reinert K, Samonte RV, et al. (2002) Recent Segmental Duplications in the Human Genome. Science 297: 1003–1007.

29. 1000 Genomes Project Human Decoy Sequences (37d5). ftp://ftp.1000genomes.ebi.ac.uk/vol1/ftp/technical/reference/phase2_reference_assembly_sequence/.

30. Benson DA, Clark K, Karsch-Mizrachi I, Lipman DJ, Ostell J, et al. (2014) GenBank. Nucleic Acids Res 42: D32–D37.

31. Levy S, Sutton G, Ng PC, Feuk L, Halpern AL, et al. (2007) The Diploid Genome Sequence of an Individual Human. PLoS Biol 5: e254.

32. Genovese G, Handsaker Robert E, Li H, Kenny Eimear E, McCarroll Steven A (2013) Mapping the Human Reference Genome s Missing Sequence by Three-Way Admixture in Latino Genomes. Am J Hum Genet 93: 411–421.

33. Tsaponina O, Haber James E (2014) Frequent Interchromosomal Template Switches during Gene Conversion in S. cerevisiae. Molecular Cell 55: 615–625.

34. Globus ST (2013) From Start to Finish: Fine Scale Mapping of Meiotic Double Strand Breaks and Gene Conversion Tracts Reveals New Insights Into Homologous Recombination: Cornell University.

35. Martini E, Borde V, Legendre M, Audic S, Regnault B, et al. (2011) Genome-Wide Analysis of Heteroduplex DNA in Mismatch Repair–Deficient Yeast Cells Reveals Novel Properties of Meiotic Recombination Pathways. PLoS Genet 7: e1002305.

36. St. Charles J, Petes TD (2013) High-Resolution Mapping of Spontaneous Mitotic Recombination Hotspots on the 1.1 Mb Arm of Yeast Chromosome IV. PLoS Genet 9: e1003434.

37. Yin Y, Petes TD (2013) Genome-Wide High-Resolution Mapping of UV-Induced Mitotic Recombination Events in Saccharomyces cerevisiae. PLoS Genet 9: e1003894.

38. Fledel-Alon A, Wilson DJ, Broman K, Wen X, Ober C, et al. (2009) Broad-Scale Recombination Patterns Underlying Proper Disjunction in Humans. PLoS Genet 5: e1000658.

39. Antonacci F, Kidd JM, Marques-Bonet T, Ventura M, Siswara P, et al. (2009) Characterization of six human disease-associated inversion polymorphisms. Human Molecular Genetics 18: 2555–2566.

40. Broman KW, Matsumoto N, Giglio S, Martin CL, Roseberry JA, et al. (2003) Common long human inversion polymorphism on chromosome 8p. In: Goldstein DR, editor. Science and Statistics: A Festschrift for Terry Speed. IMS Lecture Notes-Monograph Series. pp. 237–245.

41. Fledel-Alon A, Leffler EM, Guan Y, Stephens M, Coop G, et al. (2011) Variation in Human Recombination Rates and Its Genetic Determinants. PLoS One 6: e20321.

42. The 1000 Genomes Project Consortium (2012) An integrated map of genetic variation from 1,092 human genomes. Nature 491: 56–65.

43. Ségurel L, Wyman MJ, Przeworski M (2014) Determinants of Mutation Rate Variation in the Human Germline. Annual Review of Genomics and Human Genetics 15: 47–70.

44. Mancera E, Bourgon R, Brozzi A, Huber W, Steinmetz LM (2008) High-resolution mapping of meiotic crossovers and non-crossovers in yeast. Nature 454: 479–485.

45. Lesecque Y, Mouchiroud D, Duret L (2013) GC-Biased Gene Conversion in Yeast Is Specifically Associated with Crossovers: Molecular Mechanisms and Evolutionary Significance. Molecular Biology and Evolution 30: 1409–1419.

46. Campbell CD, Chong JX, Malig M, Ko A, Dumont BL, et al. (2012) Estimating the human mutation rate using autozygosity in a founder population. Nat Genet 44: 1277–1281.

47. Li H, Durbin R (2009) Fast and accurate short read alignment with Burrows–Wheeler transform. Bioinformatics 25: 1754–1760.

48. Patterson N, Price AL, Reich D (2006) Population Structure and Eigenanalysis. PLoS Genet 2: e190.

49. McVean GAT, Myers SR, Hunt S, Deloukas P, Bentley DR, et al. (2004) The Fine-Scale Structure of Recombination Rate Variation in the Human Genome. Science 304: 581–584.

50. Parvanov ED, Petkov PM, Paigen K (2010) Prdm9 Controls Activation of Mammalian Recombination Hotspots. Science 327: 835.

51. Hinch AG, Tandon A, Patterson N, Song Y, Rohland N, et al. (2011) The landscape of recombination in African Americans. Nature 476: 170–175.

52. Hellenthal G, Stephens M (2006) Insights into recombination from population genetic variation. Current Opinion in Genetics & Development 16: 565–572.

